# Sickle cell status skews malaria parasite genotype at infection

**DOI:** 10.1101/2025.09.09.675015

**Authors:** Helena D. Hopson, Alejandra Herbert-Mainero, Aline Gaelle Bouopda-Tuedom, Charlène Tina Nanssong-Vomo, Brigitte Tumamo Fotso, Belinda Claire Kiam, Ibrahima Ibrahima, Clément Onguene, Luc Abate, Radoslaw Igor Omelianczyk, Heather D. Evans, Lyra E. Horton, Gavin Band, Tracey J. Lamb, Lawrence S. Ayong, Sandrine E. Nsango, Ellen M. Leffler

**Author notes:** Corresponding authors: Sandrine E. Nsango, Ellen M. Leffler.

## Abstract

Sickle cell hemoglobin (HbS) confers protection against symptomatic malaria caused by *Plasmodium falciparum*. HbS carriers with symptomatic malaria harbour parasites enriched for sickle-associated alleles (*Pfsa+*), which appear to partially overcome HbS-mediated protection, but their role in asymptomatic infections is unclear. Here, we conducted a cross-sectional survey of 2,246 healthy school children in a region of high malaria transmission region in Cameroon. Analysing parasite *Pfsa* and human HbS genotypes for 1,701 asymptomatic *P. falciparum* infections confirmed that heterozygous HbS (HbAS) and homozygous non-HbS (HbAA) genotypes have similar rates of asymptomatic infection. However, the HbS genotype is strongly associated with parasites carrying *Pfsa*+ alleles at *Pfsa1* and *Pfsa3* loci, indicating a selective advantage for these alleles in HbS carriers. Our findings reveal that the protective effect of HbS is complex, where HbS has a protective effect prior to symptoms against *Pfsa-* parasites and *Pfsa+* alleles may contribute to lower rates of symptomatic disease in HbS carriers. HbS protection against malarial disease should consider both resistance and tolerance and ongoing co-evolution with parasites.

## Main Text

Sickle cell hemoglobin (HbS) is strongly protective against malaria caused by *Plasmodium falciparum*, conferring ∼90% protection against severe malaria and ∼70% protection against mild disease ^1^. However, epidemiological evidence shows that HbS either does not protect or has only a weak effect against asymptomatic infection ^1–3^. HbS has also been associated with reduced development of symptomatic malaria from asymptomatic infection ^4^. Together, these observations suggest HbS carriers tolerate *P. falciparum* infection but are protected from disease symptoms. The mechanism of HbS protection from malaria is not fully understood, but may involve restricting contributors to pathogenesis such as surface expression of parasite virulence proteins ^5^ and parasite sequestration ^6^, or by limiting immunopathology, for example by host upregulation of the stress response gene *heme oxygenase-1 (HO-1*) ^7,8^. Reduced parasite densities in HbS carriers ^9,10^ suggests the protective mechanism of HbS may also involve limitations to parasite growth due to HbS polymerization in low-oxygen conditions ^11,12^.

The effects of the HbS allele have been measured without considering genetic variation in the *P. falciparum* parasite, a unicellular eukaryotic species with a complex lifecycle alternating between human and mosquito hosts. Recently, multiple alleles in four regions of the parasite genome were found to be enriched in HbS carriers with severe and mild malaria (named *Pfsa1-4* for *Plasmodium falciparum sickle-associated*) ^13,14^. These primarily include protein coding changes: at *Pfsa1* to acyl-CoA synthetase 8 (ACS8, *PF3D7_0215300),* at *Pfsa2* to an exported protein (PF3D7_0220300), at *Pfsa3* to a putative phosphatase (PF3D7_1127000), and at *Pfsa4* to a serine/threonine kinase (FIKK4.2, PF3D7_0424700). Parasites carrying the HbS-associated alleles (*Pfsa+*) appear to partially overcome the protection conferred by HbS against malaria. Furthermore, linkage disequilibrium (LD) between *Pfsa* variants and significant differentiation across populations suggest they are under natural selection ^13–16^. However, theoretical models predict that host mutations conferring resistance from disease but not infection, i.e. disease tolerance, would not place strong selective pressure on pathogens ^17^. It has been proposed that HbS may even be favorable to *P. falciparum* parasites, with some studies reporting HbS to be associated with longer infections and higher transmission ^18,19^. Thus, it remains unclear whether HbS is driving selection at the *Pfsa* loci.

In malaria-endemic regions where HbS is prevalent, the vast majority of *P. falciparum* infections are asymptomatic ^20^, constituting most of the reservoir for transmission ^21^. We thus reasoned that evolutionary dynamics of *Pfsa* alleles may be largely determined by their fitness in asymptomatic infections. Here, we test for association between HbS and *Pfsa* alleles in asymptomatic infections by conducting a cross-sectional survey of 2,246 school children in a region of high malaria transmission in Mfou, Cameroon. We develop an amplicon sequencing panel for both host and parasite loci and obtain *Pfsa* and HbS genotypes for 1,701 asymptomatic infections. We find that heterozygous (HbAS) and homozygous non-HbS (HbAA) children have similar rates of asymptomatic infection, but that HbAS genotype is strongly associated with parasites carrying *Pfsa+* alleles. Our findings reveal a protective effect of HbS against asymptomatic infection that depends on the parasite genotype at *Pfsa* variants, demonstrating that HbS likely acts as a strong selective pressure driving co-evolution at the *Pfsa* loci.

### HbS is not protective against asymptomatic infection

We collected finger-prick blood samples from 2,685 children aged 3–17 from 11 villages in the Mfou district of Cameroon during three high malaria transmission seasons between October 2022-December 2023 (Extended Data Fig. 1). Children were enrolled while at school, where they appeared healthy and without a fever (temperature <= 37.5°C) and reported no antimalarial treatment or malaria symptoms for the past month ^22^. To genotype samples, we developed an amplicon sequencing panel with targets including the human HbS mutation (rs334, GRCh38 chr11:5227002 T>A) and parasite SNPs previously reported to be associated with HbS in severe malaria cases. Specifically, we targeted the most strongly associated SNP at *Pfsa1* (chr2: 631190 A > T, Y40F in ACS8) and *Pfsa3* (chr11: 1058035 A > T, D7V in PF3D7_1127000) ^13^, capturing additional variants within the length of each amplicon. We also genotyped four unlinked parasite loci that are known to be highly polymorphic, enabling inference of the number of parasite strains in each infection (Table S1). Following quality control, 2,246 individuals were included for analysis and genotyped in all amplicons using bcftools ^23^, with >97% genotyping success at all variants (Extended Data Fig. 2). The allele frequency of HbS was 0.09, consistent with previous surveys in the region ^18,24^, and our data included 439 heterozygous HbAS individuals (19.6%) and two homozygous HbSS individuals (0.09%). There were no significant differences in sex or age between HbAA and HbAS individuals (chi-square test p > 0.05, Table S2). Using parasite amplicon sequencing coverage, we identified 1,701 children with *P. falciparum* infections and 545 children without parasites detected (infection prevalence = 76%), again consistent with previous studies in the region ^25,26^. Parasite sequencing coverage was correlated with parasitemia categories recorded by microscopy (Extended Data Fig. 3).

There was no significant difference between HbAA and HbAS genotype groups in asexual parasite infection rate inferred either by sequencing coverage (76.4% vs. 72.9%, Wald test p = 0.16, Fig. 1A) or by microscopy (75.2% vs. 71.8%, chi-square test p = 0.16, Extended Data Fig. 4). Likewise, there was no significant difference in presence of gametocytes, the mosquito transmissible parasite form, detected by microscopy (14.1%. vs. 15.7%, Wald test p = 0.12, Fig. 1A). *P. falciparum* is haploid in the human host, such that sequence variation indicates the presence of more than one genetically distinct *P. falciparum* strain. We estimated the number of strains present in each infection (complexity of infection, COI) from haplotype variation at the polymorphic *SERA2* and *AMA1* loci using SeekDeep ^27^ and found that most infections were polyclonal (mean COI = 2.3, 71.6% and 69.9% polyclonal estimated from *SERA2* and *AMA1*, respectively), consistent with high transmission rates and previous estimates in the region ^28^. The HbAS genotype showed lower levels of polyclonality at both loci (proportion polyclonal at SERA2 = 66% HbAS vs. 73% HbAA, Wald test p = 0.012; proportion polyclonal at AMA1 = 65% HbAS vs. 71% HbAA, Wald test p = 0.048 Fig. 1B) ^4,18,29,30^ and with lower parasitemia levels (Wald test p = 0.035 Fig. 1C), as has also been found in some but not all previous reports ^4,9,10,18,31,32^. The association of HbS genotype with parasitemia and polyclonality estimated from the AMA1 amplicon were no longer significant upon inclusion of village and season as covariates (Wald test p = 0.089 and p = 0.074, respectively), but remained significant for polyclonality estimated from the SERA2 amplicon (Wald test p = 0.019).

**Fig. 1.**
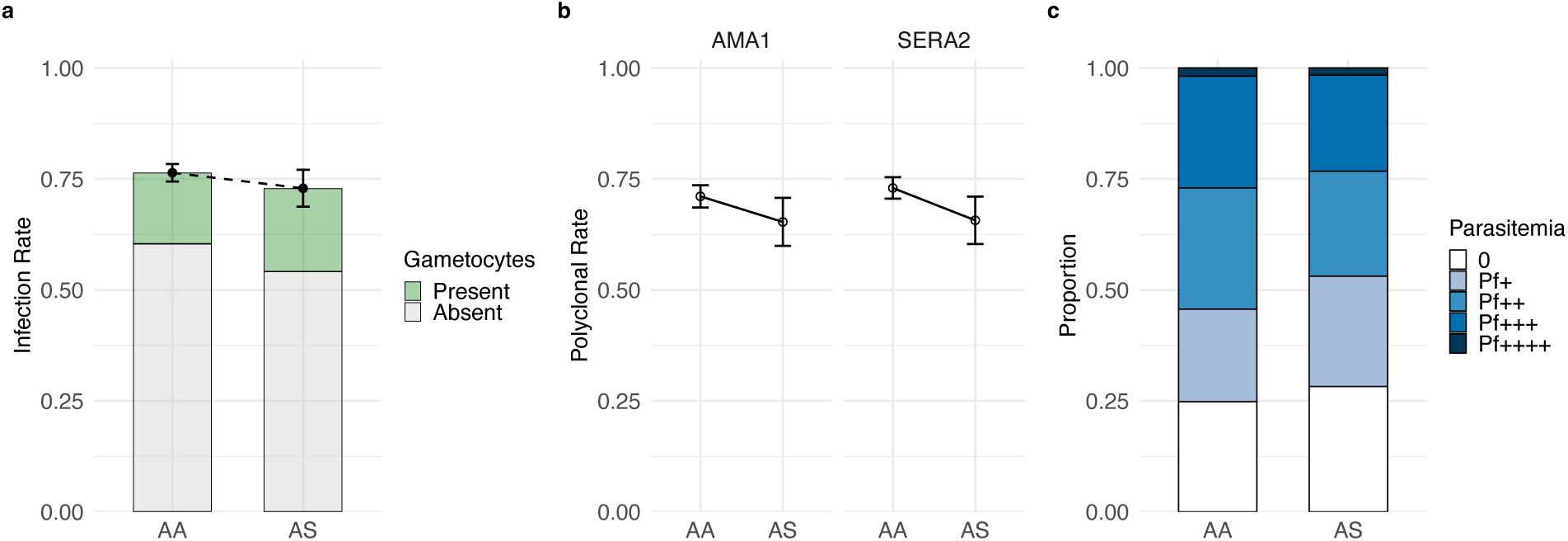
HbS is not associated with asymptomatic infection rate and is associated with lower COI and parasitemia. **(A)** The rate of asymptomatic infection determined by sequencing colored by whether gametocytes were present or absent according to microscopy (1805 HbAA and 439 HbAS individuals, chi-square test p = 0.16). Error bars represent 95% Wald confidence intervals for the unadjusted proportion of individuals infected. **(B)** Polyclonal infection rate estimated as the proportion of infections with a complexity of infection (COI) > 1 for AMA1 (n samples with COI estimate = 1549, chi-square test p = 0.06) and SERA2 (n = 1601, chi-square test p = 0.014). Error bars represent 95% Wald confidence intervals for the unadjusted proportion of individuals with polyclonal infections. **(C)** Proportion of sequencing-positive infections colored by parasitemia level from microscopy. Sample sizes: n= 1379 for HbAA and n= 320 for HbAS, Wald test p = 0.035. P-values in A and B are from two-sided Pearson’s chi-square tests and in C from a Wald test under an additive model computed using SNPTEST. No adjustments were made for multiple comparisons or covariates.

### *Pfsa1* and *Pfsa3* are strongly associated with HbS in asymptomatic infections

At individual SNPs, we interpreted homozygous genotype calls made by bcftools as indicating the presence of parasites carrying only one of the two alleles (“unmixed genotype”), and heterozygous calls with at least two reads for each allele to indicate the presence of multiple parasite strains with different alleles (“mixed genotype”). First considering unmixed genotypes at *Pfsa1* (chr2: 631190) and *Pfsa3* (chr11: 1058035), we observed a striking difference between human genotype groups, with *Pfsa+* almost exclusively in HbAS hosts and *Pfsa-* almost entirely in HbAA hosts (Fig. 2A,B). To evaluate this finding statistically, we tested for association with HbS using a logistic regression in HPTEST ^13^. These SNPs in *Pfsa1* and *Pfsa3* were strongly and significantly associated with HbS (odds ratio (OR)=60, 95% confidence interval (CI)=29-120, Bonferroni adjusted p-value (p.adj) = 2.7 × 10^-28^ for the *Pfsa1* SNP chr2:631190 T>A and OR=129, CI=55-298, p.adj = 4.9 × 10^-28^ for the *Pfsa3* SNP chr11:1058035 T>A; Table S3). No SNPs in the other amplicons were associated (n = 37, all p.adj > 0.05, Table S4). Of the five SNPs we genotyped at *Pfsa1,* the most strongly associated (chr2:631190) was also the top association in the two previous studies carried out in symptomatic malaria ^13,14^ and we refer to this variant as *Pfsa1* throughout. The association between HbS and both *Pfsa* loci was observed in all villages and seasons, and associations remained significant when including village and season as covariates (Extended Data Fig. 5 and Table S5).

**Fig. 2.**
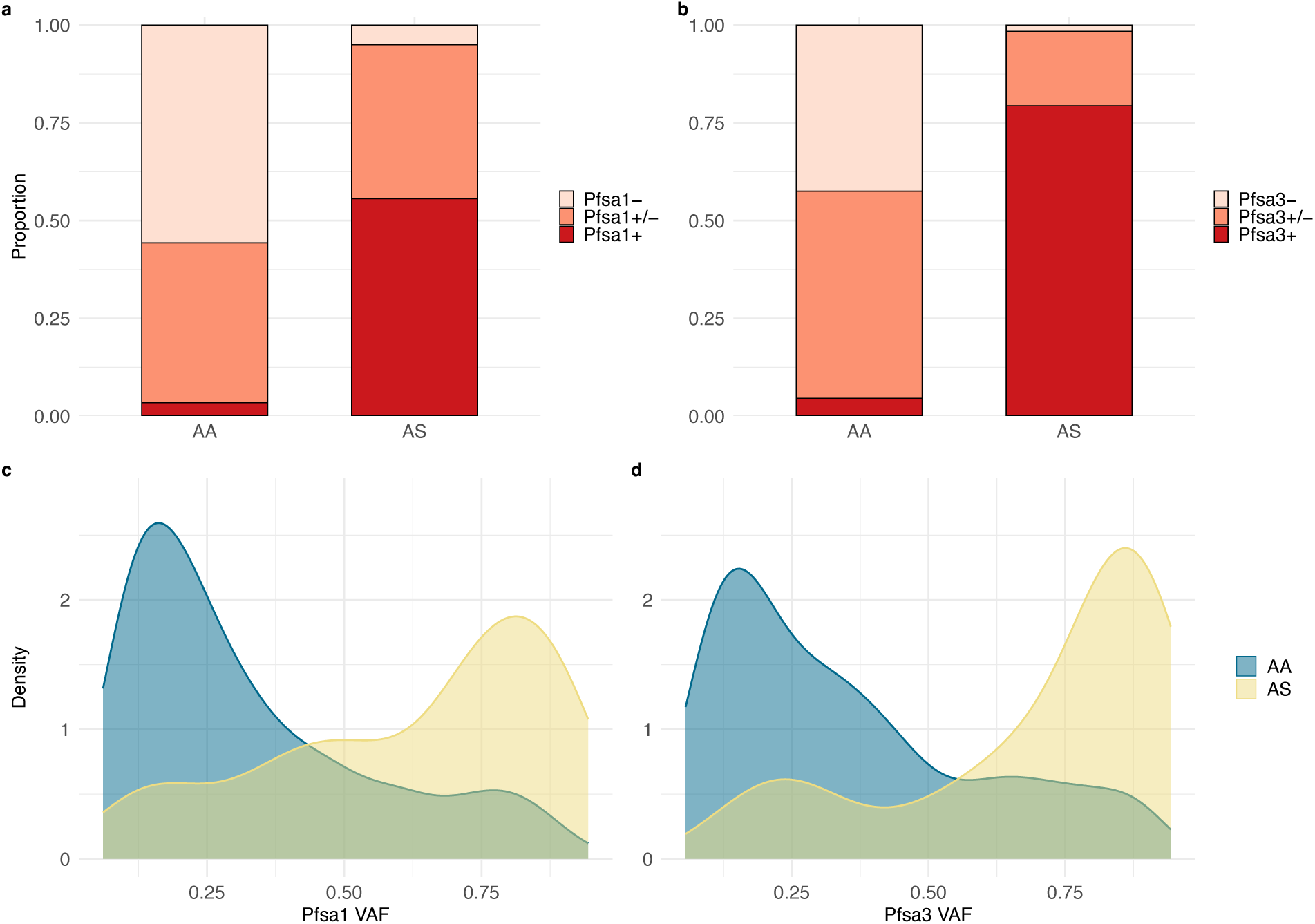
HbS is associated with *Pfsa* genotype. The proportion of asymptomatic individuals with parasites detected in their blood in HbAA and HbAS hosts, as determined from parasite sequencing reads, colored by parasite genotype for **(A)** *Pfsa*1 (chr2: 631190 within *ACS8*) and **(B)** *Pfsa*3 (chr11: 1058035 within *PF3D7_1127000*). Red are infections with parasites carrying *Pfsa*+ alleles, salmon are infections with multiple parasite strains carrying both *Pfsa*+ and *Pfsa*-alleles, and pink are infections with parasites carrying *Pfsa*-alleles. Sample sizes: n =1379 for HbAA and n =320 for HbAS in (A), and in (B) n = 1346 for HbAA and n =310 for HbAS. Comparing unmixed genotype counts (n = 1009) for HbAA vs HbAS, p = 2.3 × 10^-146^ in (A), and in (B) unmixed genotype counts (n = 884) for HbAA vs HbAS, p = 7.8 × 10^-139^. P-values in A and B are from a two-sided Pearson chi-square test. **(C)** *Pfsa1* variant allele fraction within individual infections containing both *Pfsa*1+ and *Pfsa*1-carrying parasite strains in HbAA (blue) and HbAS (yellow). Sample sizes: n = 569 for HbAA and n = 130 for HbAS, p = 2.2 × 10^-25^. **(D)** *Pfsa3* variant allele fraction within individual infections containing both *Pfsa*3+ and *Pfsa*3-carrying parasite strains. Sample sizes: n = 713 for HbAA and n = 59 for HbAS, p = 3.4 × 10^-17^. P-values in C and D are from a two-sided Wilcoxon rank-sum test.

Across all infections, we observed 40.6% (690/1701) with mixed genotype at *Pfsa1* (Fig. 2A) and 46.6% (772/1657) with mixed genotype at *Pfsa3* (Fig. 2B). The high number of mixed genotype infections, common in asymptomatic infections in high transmission regions ^25,33^, allowed us to test for competition between *Pfsa+* and *Pfsa-* parasites within individual infections by looking for skewed allelic composition. We found that the variant allele fraction (VAF) was highly skewed towards parasites carrying *Pfsa-* alleles in HbAA hosts and towards parasites carrying *Pfsa*+ alleles in HbAS hosts for both *Pfsa1* (Fig. 2C) and *Pfsa3* (Fig. 2D), and this pattern remained at more stringent coverage depth requirements (Extended Data Fig. 6).

These results across and within infections indicate that HbS confers substantial resistance to blood-stage infection by *Pfsa-* parasites, whereas *Pfsa+* parasites have a selective advantage. Consequently, *Pfsa* alleles not only confound estimates of HbS protection against disease ^13,14^ but also asymptomatic infection. These findings revise the interpretation that *Pfsa+* parasites overcome HbS protection against disease ^34,35^, instead indicating that *Pfsa+* parasites commonly infect HbS hosts but few overcome HbS disease protection.

To indirectly assess if *Pfsa* alleles influence the conversion rate from asymptomatic infection to symptomatic disease, we compared the effect sizes of the HbS-*Pfsa* association estimated here in asymptomatic infections with those previously reported in symptomatic malaria cases. Effect sizes are larger in asymptomatic infections for both *Pfsa1* and *Pfsa3* (Fig. 3A). This difference could be explained if *Pfsa-* parasites are more likely to cause symptoms in HbS carriers, or if *Pfsa+* parasites are more likely to cause symptoms in HbAA individuals, as either would reduce the effect size of the HbS/*Pfsa* association in symptomatic malaria relative to asymptomatic. Comparing across studies, we observe a lower proportion of *Pfsa+* infections in asymptomatic HbAA hosts than has been reported in HbAA hosts with severe malaria (Fig. 3B). Although interpretation is limited because the populations differ in HbS and *Pfsa* allele frequency and transmission intensity among other factors, this observation suggests that parasites carrying *Pfsa+* alleles may be more virulent than parasites with *Pfsa-* alleles in HbAA hosts. We identified a study in the Gambia ^36,37^ where higher frequencies of *Pfsa+* were found in symptomatic relative to asymptomatic infections, consistent with this prediction (Table S6). Further studies that directly compare asymptomatic and symptomatic infections in the same population and/or longitudinally will help clarify this potential relationship.

**Fig. 3.**
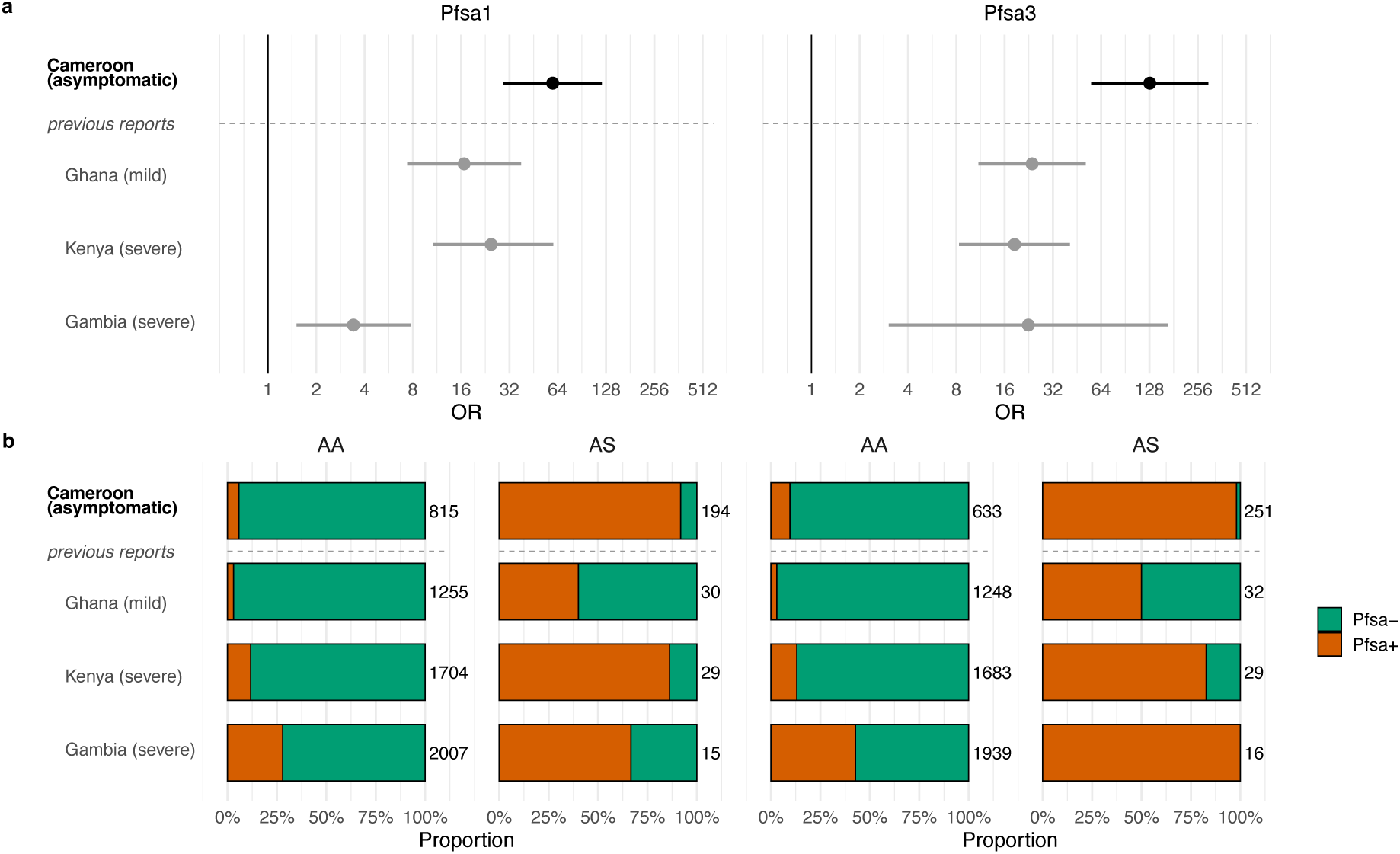
Strength of HbS-*Pfsa* association across populations and disease severity. **(A)** Comparison of effect sizes for the association of HbS and *Pfsa*+ alleles (left – *Pfsa*1+ (chr2: 631190), right – *Pfsa*3+ (chr11: 1058035)) from this study (bold) with previously reported effect sizes in symptomatic malaria (light grey). Number of HbAA/HbAS/HbSS individuals included in association tests: Cameroon - 815/194/2 (*Pfsa1*) and 633/251/1 (*Pfsa3*), Ghana - 1255/30/1 (*Pfsa1*) and 1248/32/1 (*Pfsa3*), Kenya - 1704/29/8 (*Pfsa1*) and 1683/29/7 (*Pfsa3*), Gambia - 2007/15/3 (*Pfsa1*) and 1939/16/3 (*Pfsa3*). Error bars represent 95% confidence intervals for the odds ratio estimates from logistic regression. **(B)** Proportion of infections in HbAA and HbAS hosts with parasites carrying *Pfsa*+ (orange) and *Pfsa*- (green) alleles and sample sizes from the same studies as in (A). HbSS individuals were included in the logistic regression and odds ratio estimates but excluded from the plots because there were very few. Infections with mixed *Pfsa* genotypes were not included in association testing in any of the studies. The Ghana data are from ^14^ and Kenya and Gambia data are from ^13^.

### *Pfsa1* and *Pfsa3* allele co-occurrence suggests epistasis

Across all infections with unmixed genotypes (HbAA = 528; HbAS = 171), *Pfsa1* and *Pfsa3* variants were in strong inter-chromosomal linkage disequilibrium (LD) (r^2^=0.84, p < 2 × 10^-16^) driven by overrepresentation of parasites carrying neither or both + alleles (*Pfsa1-/Pfsa3-* and *Pfsa1+/Pfsa3+*) and underrepresentation of parasites carrying one + allele but not the other (*Pfsa1-*/*Pfsa3+* and *Pfsa1+*/*Pfsa3-*) (Fig. 4A, Table S7). This pattern of inter-chromosomal LD was unique to *Pfsa1* and *Pfsa3* and absent across all other pairs of *P. falciparum* SNPs genotyped in this dataset (Table S8), agreeing with previous observations genome-wide ^13^. LD was lower but remained statistically significant when assessed separately in HbAA and HbAS infections (r^2^=0.59, p < 2 × 10^-16^ in HbAA and r^2^=0.26, p = 1 × 10^-12^ in HbAS). Compared with expectations in the absence of LD, in HbAA the small number of parasites carrying a + allele at one *Pfsa* but not the other was particularly striking (10- and 20-fold lower than expected for *Pfsa1-/Pfsa3+* (n = 12) and *Pfsa1+/Pfsa3-* (n = 5), respectively, two-sided binomial test p < 1 × 10^-5^; Fig. 4A, Table S7). In contrast, *Pfsa1+*/*Pfsa3+* parasites (n = 31) were only 1.5-fold lower than expected (two-sided binomial test p = 0.03), suggesting an epistatic effect whereby the *Pfsa+* alleles have a higher fitness cost when carried with *Pfsa-* alleles. In HbAS infections, the majority of infections (94%) had *Pfsa1+*/*Pfsa3+* parasites (n = 160) which was 11-fold more than expected (two-sided binomial test p < 1 × 10^-5^). All other combinations were significantly underrepresented, with *Pfsa1-/Pfsa3-* (n = 3) showing the largest difference from expected (28-fold lower; two-sided binomial test p < 1 × 10^-5^ Fig 4A, Table S7).

**Fig. 4.**
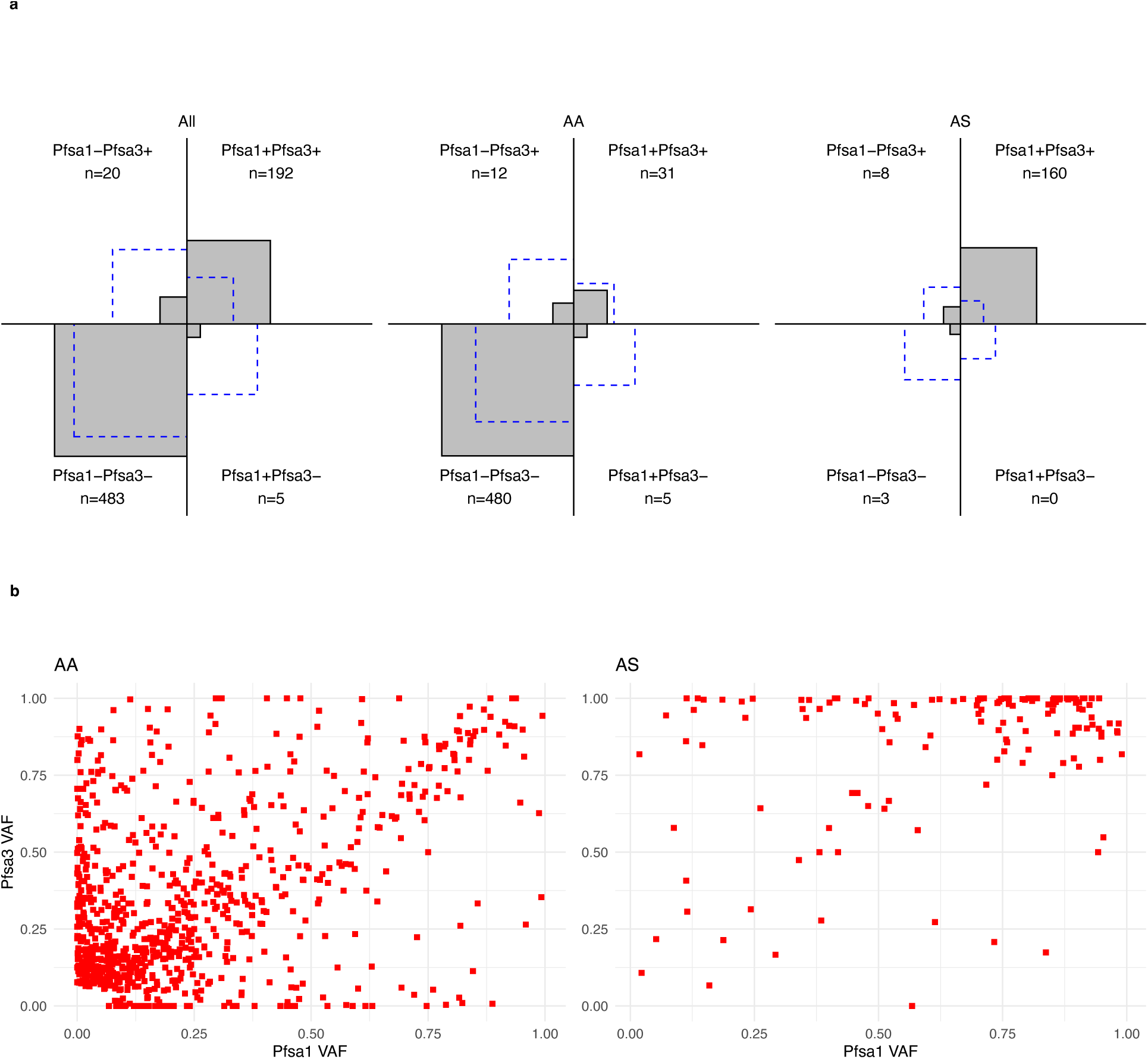
*Pfsa*1 and *Pfsa*3 are in strong inter-chromosomal linkage disequilibrium. **(A)** Comparison of observed and expected numbers of genotype combinations between *Pfsa*1 and *Pfsa*3. The area of each square is proportional to the number of infections with a particular combination of alleles with the observed number in gray and value provided, while the expected number is shown in blue, calculated using the *Pfsa*1 and *Pfsa*3 allele frequencies (0.28 and 0.3) in all samples non-mixed at both *Pfsa*1 and *Pfsa*3 (n = 700). Expected and observed counts are shown for all individuals as well as separately for HbAA (n=528) and HbAS (n=171). **(B)** VAF for *Pfsa*1 and *Pfsa*3 in infections with mixed genotypes at either *Pfsa*1 and *Pfsa*3, separately for HbAA and HbAS.

Among 983 infections with a mixed genotype at either *Pfsa1* or *Pfsa3*, we observed a correlation in VAF between *Pfsa1+* and *Pfsa3+* alleles in both HbAA and HbAS infections indicating LD within infections (r^2^=0.21, p < 2 × 10^-16^ and r^2^=0.14, p = 4.5 × 10^-6^, respectively, Fig. 4B). This correlation is lower than in infections with unmixed *Pfsa* genotypes (Fig 4A), and we also see evidence of strains carrying one *Pfsa+* allele and one *Pfsa-* allele at the two loci, indicated by points off the diagonal (Fig 4B). Many of these mixed genotype infections likely derive from co-transmission of *Pfsa1+/Pfsa3+* and *Pfsa1-/Pfsa3-* parasites by a single mosquito in which meiosis occurred generating *Pfsa1*+/*Pfsa*3- and *Pfsa1*-/*Pfsa*3+ parasites. Although some of the LD may be re-established in the mosquito or liver stages, the observation that *Pfsa1+/Pfsa3-* and *Pfsa1-/Pfsa3*+ strains are present in mixed infections indicates competition during blood-stage infection contributes to maintain LD. Future studies that can distinguish between superinfection (different clones introduced from independent mosquitoes) and co-transmission (recombination between different clones in one mosquito) by incorporating parasite relatedness, as well as the age of infection, will clarify the role of intra-host competition in selection for *Pfsa* allele combinations.

## Discussion

In our study of 2,246 healthy children in the Mfou region of Cameroon, we uncovered a strong association between human HbS genotype and *P. falciparum Pfsa1* and *Pfsa3* genotype in asymptomatic infections. We found no significant difference in the rate of asymptomatic infection between HbAA and HbAS children, but that both *Pfsa+* alleles are at higher frequency in HbAS hosts than in HbAA hosts, consistent with a selective advantage of *Pfsa+* alleles in HbS carriers at infection. LD patterns between the parasite loci suggest there is selection on *Pfsa* alleles in HbAA hosts as well and that it may involve epistasis. The near absence of *Pfsa-* alleles in HbS hosts reveals that HbS confers resistance to *Pfsa-* parasites separate from a tolerance effect that protects against disease symptoms from *Pfsa+* parasites. Our results highlight that the protective mechanism of HbS is complex, involving both resistance to infection and tolerance to disease, and requires consideration of *Pfsa* genotype when teasing out underlying mechanisms. In this study, we have focused on *Pfsa1* and *Pfsa3*, which are present in parasites across sub-Saharan Africa with frequency correlated to that of HbS. Because previously reported sickle associated variants in the *Pfsa2* locus are absent in Central and West Africa including Cameroon, it was excluded from our study ^13,38^. We could not confidently genotype the *Pfsa4* variants by this approach due to nearby repetitive sequence. Future incorporation of *Pfsa2* and *Pfsa4* genotypes in relevant populations could yield insight into their role and potential epistatic interactions between these additional loci. All four genes are not well characterized or overtly functionally related (functional evidence summarized here^13,14^). Our work suggests that they are likely to have functions important for establishment of infection preceding disease symptoms.

The observed lack of protection conferred by HbS against asymptomatic infection is consistent with many but not all previous reports ^1,39,40^. Our results suggest that whether a protective effect by HbS is observed in asymptomatic infections could depend on factors including the transmission intensity, acquired antimalarial immunity, and the frequency of *Pfsa+* alleles in the population. In a high transmission setting such as this study, where children are frequently exposed to both *Pfsa+* and *Pfsa-* parasites, equal infection rates in HbS and non-HbS are driven by different underlying parasite genotypes. In contrast, in a low transmission setting, where children are exposed more often to the more common *Pfsa* allele, there may be a higher asymptomatic infection rate in HbAA or HbAS children, depending on which *Pfsa* allele is more common. Future studies of HbS-*Pfsa* association in asymptomatic infection across a range of transmission and allele frequencies will further elucidate the dynamics of this system and whether these factors explain some of the variability in previous reports that only considered HbS genotypes. It is plausible that other host factors impact the interaction between HbS and parasite genotypes. For example, variation in fetal hemoglobin (HbF) levels has been reported and linked to sickle cell disease severity ^41^. Future work incorporating genetic variants associated with HbF or directly measuring HbF levels could uncover such interactions.

Our findings suggest a revised evolutionary model for HbS protection against malaria that involves ongoing co-evolution with *P. falciparum* parasites. A plausible model is that when HbS arose an estimated 10,000-50,000 years ago ^42^, it conferred resistance to all *P. falciparum* parasites (*Pfsa-*), which in turn selected for *Pfsa+* parasites. This pattern reflects co-evolutionary dynamics whereby hosts evolve resistance, which selects for counteradaptation in parasites, which then selects for enhanced host resistance, etc. resulting in non-equilibrium co-evolutionary dynamics ^43^. However, the equal infection rates in HbAA and HbAS driven by *Pfsa-* and *Pfsa+* parasites, respectively, and previously observed correlated frequencies of HbS and *Pfsa* alleles suggest a possible stable equilibrium state ^13^. The HbS allele is under balancing selection and maintained at an intermediate frequency by the advantage of protection against malaria and the fitness cost of sickle cell disease in homozygous individuals (HbSS) ^44,45^. It is plausible that *Pfsa+* alleles are in turn maintained under balancing selection by the advantage of infecting HbS hosts and a fitness cost in HbAA hosts. Haplotype patterns and geographical co-distribution of alleles offer additional support for this hypothesis ^46^. The functional mechanisms, fitness tradeoffs and long-term temporal dynamics of these and the other *Pfsa* alleles remain unclear and will be important for future studies to uncover.

## Methods

### Ethics and consent

This study was approved by the National Ethics Committee for Human Health Research in Cameroon under agreement 2021/06/1366/CE/CNERSH/SP. All participants were enrolled as volunteers after their legal parent/guardian signed the informed consent form. Blood samples were used exclusively as de-identified samples to safeguard donors’ privacy. Confidentiality was maintained and no identifying information was disclosed or published without the donor’s consent.

### Study site and samples

A cross-sectional survey was performed in the Mfou district of Cameroon over three seasons of high malaria transmission (October-December 2022, April-May 2023, and November-December 2023). Children in school who met the following study criteria were enrolled: parental consent, no fever, and self-reported no malaria infection or anti-malarials taken in the past month. For each individual, blood was collected from finger pricks onto Whatman Grade 17 paper (dried blood spots, DBS) and a microscope slide. The presence of asexual parasites and gametocytes was assessed by qualified microscopists from microscopic examination of Giemsa-stained blood smears using a Leica® light microscope with a 100× oil-immersion objective. Semi-quantification of parasitemia was categorically grouped into levels from 0-5, corresponding to the number of *P. falciparum* trophozoites per µl of blood as follows: 0 = absence of trophozoites / field crossing (0 trophozoites/µl), Pf+ = 1-10 trophozoites / crossing (1-50 trophozoites/µl), Pf++ = 10-20 trophozoites / crossing (51 – 500 trophozoites/µl), Pf+++ = 1-10 trophozoites / field (501 – 5,000 trophozoites/µl), Pf++++ = 11-100 trophozoites / field (5,001 – 50,0000 trophozoites/µl), Pf+++++ = over 100 trophozoites / field (> 50,000 trophozoites/µl) ^47^. Sex, age, and temperature were also recorded. Parasite positive children received artemisinin-based combination therapy (ACT) in accordance with national guidelines. Metadata for each participant is provided in Table S9.

### Amplicon sequencing

Genomic DNA was extracted from dried blood spots by Autogen (https://autogen.com/) on the XTRACT 16+ automated magnetic bead DNA isolation platform. We developed both a human and parasite amplicon panel for targeted sequencing of select loci. The human panel included amplicons covering the HbS variant and other variants associated with protection from severe malaria (in *ABO, G6PD,* and *ATP2B4*) ^24^, which will be presented elsewhere. Primers for human amplicons were purchased from Thermo Fisher Scientific. The parasite panel included newly designed amplicons covering sickle-associated variants in *Pfsa1*, *Pfsa3*, and *Pfsa4*, as well as amplicons targeting the highly polymorphic parasite genes *SERA2, AMA1, CSP,* and *TRAP.* Primer sequences for *SERA2, AMA1, CSP,* and *TRAP* were adopted from the previously described 4CAST amplicon panel ^48^ and primers targeting the *Pfsa* loci were designed for this study using the oligonucleotide design tools in Benchling. Primers were evaluated for target specificity using BLAST and assessed for compatibility within the multiplex primer pool using online tools that evaluate factors including dimer formation and melting temperatures. Primer sequences, genomic coordinates, and references and catalog numbers are provided in Table S1. *Pfsa2* is not polymorphic in Cameroon and was not included.

We modified the published 4CAST amplicon sequencing protocol to accommodate inclusion of the additional amplicons as described below. Primers were pooled separately by species, adjusting the concentration of each primer to improve evenness of coverage across the amplicons after finding that some amplicons performed better than others. The concentration of the forward and reverse primers for each amplicon within the parasite and human primer pools are listed in Table S10 and S11. For each sample, a PCR reaction was performed separately for the human and parasite primer pool such that each sample had two PCR1 products – one with human amplicons and one with parasite amplicons. Inputs to PCR1 reaction are in Table S12.

PCR1 cycle for human amplicon panel – 1. 95.0° - 03:00, 2. (98° -00:20, 63° - 00:15, 72° - 00:30) × 25 cycles, 3. 72° - 01:00, 4. 4 ° - ∞

PCR1 cycle for parasite amplicon panel - 1. 95.0° - 03:00, 2. (98° -00:20, 57° - 00:15, 62° - 00:30) × 25 cycles, 3. 72° - 01:00, 4. 4 ° - ∞

To multiplex samples for sequencing, unique dual indexes were added to the PCR1 product in a second PCR. We used sets A-D of the Illumina DNA/RNA Unique Dual Indexes (Product numbers 20027213, 20027214, 20042666, 20042667), allowing for unique labeling of four 96 well plates (up to 384 samples). Inputs to PCR2 reaction are in Table S13.

PCR2 cycle - 95 degrees - 01:00, 2. (95 degrees - 00:15, 55 degrees - 00:15, 72 degrees 00:30) × 10 cycles, 3. 72 degrees - 01:00, 4. 4 degrees infinity

PCR2 products from a single plate were combined by adding 8 µl from each well into a single microcentrifuge tube, resulting in one human and one parasite tube of PCR2 products from 96 samples. Each tube of combined PCR2 products was purified by three rounds of AMPure XP bead clean up (Product number A63880) at a bead ratio of 0.9X each round. The concentration of each tube of combined PCR2 products was measured with Qubit BS DNA Assay (Product number Q32853). To make a final library for sequencing, we combined human and parasite plates A-D such that each plate was at the same concentration relative to the other plates with parasite products at a 2X concentration relative to the human products in order to (i) increase coverage at parasite amplicons as there were a greater number of parasite amplicons (8) compared to human amplicons (6) and (ii) increase sensitivity for lower parasitemia which is common in asymptomatic infections. Final libraries were sequenced on MiSeq 250 Cycle Paired End Sequencing v2 at the Huntsman Cancer Institute High Throughput Genomics Core at the University of Utah.

### Mapping and quality control

Following demultiplexing to sample level fastq files, Illumina adapters were trimmed with cutadapt (version 3.5) ^49^. Paired sequence reads were then mapped with bwa mem (bwa version 0.7.17) ^50^ to a joint human and parasite reference genome generated by concatenating the human GRCh38 and PlasmoDB-62 *P. falciparum* 3D7 (version=2020-09-01) reference genomes. Mean coverage of each amplicon in each sample was calculated using samtools mpileup (samtools version 1.16) ^23^ and custom scripts. The recently reported *Pfsa4* allele is flanked by repetitive regions which interfered with mapping, so this amplicon was removed from further analysis. We assigned samples as infected with *P. falciparum* parasites when they had mean sequence coverage >50 at all three parasite amplicons that had the highest coverage across all samples (AMA1, SERA2, and ACS8_1). One plate of samples was excluded because parasite sequences were recovered from negative controls (n=376). Samples without microscopy data (n = 22) were also excluded (Extended Data Fig. 2).

### Genotyping

All samples were genotyped for human amplicons and only infected samples were genotyped for parasite amplicons using bcftools mpileup and call -m with default options except increasing -- max-depth to 1000000 (bcftools version 1.16) ^23^. Variants were filtered to keep biallelic sites with a minor allele frequency greater than 0.05, quality score greater than 20, and less than 25% of missing genotypes. Genotypes with total read depth less than 10 were set to missing. This excluded 10 samples with HbS read depth < 10 (Extended Data Fig. 2). Of the remaining samples, >99% had over 30X coverage at HbS. For the parasite, which is haploid in the human host, a 0/1 genotype call indicates a mixed genotype, meaning more than one parasite clone is present and they carry different alleles at that SNP. To increase confidence in mixed genotype calls, we required that each allele was present in at least two reads. If an allele was only present in one read, the genotype was changed to a homozygous call for the other allele. To ensure that differences in VAF were not affected by sequencing errors at low read depths, we also considered thresholds of alternate and reference reads in mixed infections up to 30X (Extended Data Fig. 6).

### Complexity of infection

The 4CAST amplicon sequences (SERA2, AMA1, CSP, and TRAP) were additionally analyzed using SeekDeep v3.0.1 to infer complexity of infection from the observed haplotypes ^27^. We followed the workflow for Illumina paired-end reads without replicates and adjusted parameters to account for low density infections according to suggestions by developers and another study ^51^. We report the complexity of infection estimates from SERA2 and AMA1 because they had the highest coverage, and AMA1 had the greatest number of polymorphic sites within these samples.

### Genetic association tests

Association between HbS genotype and each phenotype was tested using SNPTEST v2.5.2 ^52^ under a frequentist additive model, with and without including village and season as categorical covariates and significance was assessed using two-sided Wald tests. Infection status and gametocyte carriage were analyzed as binary phenotypes using logistic regression. Infection status was determined from sequencing coverage as described under “Mapping and quality control” and gametocyte carriage was assessed by microscopy. Complexity of Infection (COI) and parasitemia were analyzed as continuous phenotypes using linear regression. COI ranged from 0 to 7 (inferred from AMA1 or SERA2 by SeekDeep) and parasitemia levels ranged from 0 to 4 as follows: 0 (submicroscopic), Pf+, Pf++, Pf+++, and Pf++++ or higher.

Genetic associations between HbS and parasite alleles were tested using the program HPTEST v2.1.9 ^13^, which implements a logistic regression model with host genotype as the predictor and parasite genotype as the outcome. We estimated the effect of the host allele on parasite allele under an additive model. HPTEST excludes missing and heterozygous genotypes. HPTEST outputs summary statistics for each host and parasite allele combination including a Bayes factor which describes the strength of association, P-value, effect size, and standard error. Odds ratios and confidence intervals were calculated from the effect size and standard error. We report the results of HPTEST without any covariates in the main text, but we also implemented the model with covariates village and season to evaluate possible contribution of batch effects or population structure. *Pfsa* variants are significantly associated with HbS in all models, and no other *P. falciparum* variants are associated with HbS. The summary statistics with sample and genotype count for each parasite allele tested against HbS are provided in Table S3-S5.

### Allele frequencies and linkage disequilibrium

Allele frequencies and inter-chromosomal linkage disequilibrium (LD) between *Pfsa1* and *Pfsa3* were estimated using the 700 samples with non-mixed genotype calls at *Pfsa1* and *Pfsa3*. Inter-chromosomal LD was calculated as the square of the Pearson correlation between all pairs of SNPs on different chromosomes (Table S8). To calculate the expected number of *Pfsa1* and *Pfsa3* allele combinations under a scenario in which the alleles were independently segregating and not in linkage disequilibrium, we multiplied the *Pfsa* allele frequencies for each of the four possible combinations of *Pfsa* alleles (*Pfsa1+/Pfsa3*+, *Pfsa1-/Pfsa3+, Pfsa1+/Pfsa3-, Pfsa1-/Pfsa1-)* by the total number of samples with non-mixed genotypes at *Pfsa1* and *Pfsa3* (Table S7). LD and *Pfsa* allele combinations were analyzed in the total sample and separately in HbAA and HbAS hosts.

The variant allele fraction (VAF) was calculated for each sample by dividing the number of alternate allele (*Pfsa+*) reads by the sum of reference and alternate allele reads at a particular *Pfsa* variant. A VAF closer to one indicates an infection with more *Pfsa+* reads and a VAF closer to zero indicates more *Pfsa-* reads.

### Statistics

Unless otherwise specified, statistical analyses were performed in R version 4.4.1 and where indicated, p-values were adjusted for multiple testing using a Bonferroni correction. Differences in participant characteristics were evaluated using Pearson’s chi-square tests and age was categorized into three groups (Table S2). Differences in proportions of infected individuals and polyclonal infections by HbS genotype were statistically evaluated by Pearson’s chi-square tests. Variant allele fractions in mixed *Pfsa* infections were compared between HbAA and HbAS individuals using a two-sided Wilcoxon rank-sum test. Linkage disequilibrium was quantified as the squared Pearson correlation coefficient. Pearson correlations and associated p-values were calculated using the cor.test function in R. P-values were Bonferroni adjusted for the 681 inter-chromosomal SNP pair comparisons. Expected counts of the four possible *Pfsa1*-*Pfsa3* allele combinations were calculated by multiplying their allele frequencies by number of individuals in each group (total individuals with unmixed *Pfsa1* and *Pfsa3* genotypes and stratified by HbAA and HbAS). For each allele combination, observed counts were compared with expected counts using two-sided exact binomial tests. P-values for the HPTEST analysis were Bonferroni-adjusted for 43 parasite variants tested, and associations with adjusted p-values below 0.05 were considered statistically significant.

## Ethics and Inclusion Statement

This study was designed by researchers at Centre Pasteur du Cameroun and the University of Utah, with local researchers involved throughout the research process, including study design, implementation, data collection, interpretation, and authorship. The research questions and study procedures were developed jointly by the collaborating institutions and were informed by the local epidemiological context. Participants were recruited according to predefined eligibility criteria appropriate to the local study population and research objectives. The study was reviewed and approved by the relevant ethics committees in Cameroon. This collaborative approach supported active participation of local researchers and strengthened local research capacity while ensuring that the study addressed locally relevant scientific and public health priorities.

## Data Availability

Mapped human and *P. falciparum* amplicon sequence reads that support the findings of this study have been deposited in NCBI SRA under BioProject PRJNA1503656.

## Code Availability

Source code for HPTEST is available at https://enkre.net/code/ and Zenodo (https://doi.org/10.5281/zenodo.5685581). The SNPTEST software is available at: https://www.chg.ox.ac.uk/~gav/snptest/. The code for figures and analysis is available on GitHub at: https://github.com/hopsonhelena/pfsa_asymptomatic/.

## Acknowledgements

The support and resources from the Center for High Performance Computing at the University of Utah are gratefully acknowledged. We acknowledge support from the HSC DNA Sequencing Core at the University of Utah as well as the High-Throughput Genomics Shared Resource and HCI’s Cancer Center Support Grant. We thank Antoine Claessens (Université de Montpellier) for help finding symptomatic status in the Gambian study and Paul Sigala (University of Utah) for comments on an earlier version of this manuscript. We thank the students from the visited schools, as well as their parents or guardians for their involvement. Our gratitude also goes to the teachers, school principals and the local administrative authorities for their approval and support in carrying out this project.

## Author Contributions Statement

Conceptualization: SEN, EML

Data curation: HDH, CTNV, BTF, AGBT, BCK, II, CO, LA

Formal analysis: HDH, AHM, RIO, HDE, LEH

Funding acquisition: SEN, EML

Investigation: HDH, AHM, CTNV, BTF, AGBT, BCK, II, CO, LA

Methodology: HDH, EML

Project administration: TJL, LSA, SEN, EML

Resources: LSA, SEN, EML

Supervision: SEN, EML Visualization: HDH

Writing – original draft: HDH, EML

Writing – review and editing: HDH, AHM, RIO, GB, SEN, EML

